# Climate-driven habitat loss and natural fragmentation increase extinction risk and compromise population viability in freshwater fish

**DOI:** 10.1101/2025.03.02.641059

**Authors:** Jessica Côte, Gaël Grenouillet, Fanny Thiéry, Céline Jézéquel, Bernard Hugueny

## Abstract

Climate change is one of the major drivers of the current extinction crisis. In this context, determining the probability of species extinction is crucial. In aquatic ecosystems, basin area is a key factor influencing extinction risk. Habitat fragmentation can significantly reduce river basin size and lead to both deterministic and stochastic extinctions. Deterministic extinctions occur when climatic conditions become unfavorable across the entire area of occupancy of populations, while stochastic extinctions result from temporal variability in population size driven by stochasticity events.

In this study, we developed an extinction model incorporating species presence probability, basin area, and biological parameters. Using species distribution modeling to estimate current and future species distributions, we applied the extinction model to freshwater fish species in France to quantify deterministic and stochastic extinctions as well as population viability under future climatic conditions. Our results showed a significant effect of climate change on deterministic extinction rates, depending on the species considered and their spawning thermal preferences. We also observed spatial patterns in extinction rates, with small coastal rivers being particularly impacted. Furthermore, our results showed an effect of climate change on population viability, especially for cold-water species.

While our results did not indicate high extinction rates for most species, they revealed that cold-water species could be severely affected by habitat loss due to climate change and natural fragmentation, with potentially greater impacts under anthropogenic fragmentation. The application of this model could be highly valuable for identifying both priority areas and species of concern in conservation planning.

## Introduction

The world is entering a major extinction crisis (Barnosky et al., 2011; Brook et al., 2008; S. Pimm et al., 1995; Thomas et al., 2004), with global change such as climate change, habitat loss, and invasive species being key drivers of current biodiversity loss (Ewers & Didham, 2006; Sax & Gaines, 2003). Human activities led to a hundredfold increase in current background extinction rates, and this could escalate to a thousand fold in the coming decades (S. L. Pimm, 2009). Climate change acts as a trigger for increasing extinction risk by altering species spatio-temporal distributions and leading to population declines (Thomas et al., 2004). Species can respond to new climatic conditions in two main ways: (i) through local adaptations to the new environment, or (ii) by dispersing to track spatial shifts in suitable habitat. Therefore, where and when climatically favorable habitats emerge and whether they are colonized, possibly including evolutionary processes related to funding events, are crucial factors in assessing species vulnerability to climate change (Cobben et al., 2012). In particular, the long-term persistence of a species could be compromised by the accumulation of climate-driven local extinctions that are not balanced by the colonization of newly suitable areas. While this issue is frequently recognized in the literature by using no-dispersal scenarios in Species Distribution Models (e. g. Thomas et al., 2004), very few studies have addressed dynamic climatic landscapes in combination with dispersal and local extinctions (e. g. (Talluto et al., 2017)). The primary reason for that is the lack of empirical data needed to parameterize sub-models for extinction and dispersal rates for most species. This task becomes more feasible in systems where physical barriers make dispersal among populations negligible over ecological time-frames, as observed in certain insular or fragmented systems, which we will refer to as isolates for simplicity. A pertinent example comes from boreal mammal communities inhabiting mountaintops surrounded by unsuitable xeric habitats, where the consequences of global warming were studied by McDonald & Brown, 1992. In such cases, rising temperature lead to the contraction of boreal habitats, triggering increased population extinction rates. For these and other comparable sets of isolates, McDonald & Brown, 1992 advocated using the species-area relationship (SAR) to predict the number of population extinctions due to climate-driven habitat shrinkage. The challenge with using SAR in this context is that it does not predict how long it will take for extinctions to occur (Halley et al., 2016; Hugueny, 1989a; Wearn et al., 2012), or which species are at higher risk of extinction. This creates challenges for planning and prioritizing conservation actions. What is needed is an extinction-area relationship (EAR) that predicts the lifespan of a population based on the area of suitable habitat it occupies. Such an approach has already been followed to assess the consequences of present habitat fragmentation (Broekman et al., 2022; Williams et al., 2022), but, to the best of our knowledge, no explicit quantification of population extinction rates has been applied to predict the effects of future climate change on species within a collection of isolates. The main objective of this study is to introduce such an approach in river basins, which function as ecological islands for strictly freshwater organisms (Hugueny, 1989a). Specifically, we will focus on strictly freshwater fish species confined to river basins that are isolated from each other by natural barriers (saltwater downstream and terrestrial habitats otherwise).

Freshwaters occupy less than one percent of the Earth’s surface and are naturally fragmented into numerous watersheds, making freshwater organisms more vulnerable to human pressures than their marine or terrestrial counterparts. Moreover, freshwater ecosystems are strongly exposed to human activities such as pollution, water flow regulation, overexploitation, habitat degradation, and the spread of exotic species (Dudgeon et al., 2006; Fagan, 2002; Giam et al., 2012; Nilsson et al., 2005). As a result, freshwater species are already among the most threatened on Earth (Ricciardi & Rasmussen, 1999; Urban, 2024), and the ongoing anthropogenic climate change is likely to worsen this worrying situation for many species. In particular, large and strictly freshwater organisms like fishes are likely to suffer greatly if climate change induces habitat loss. A meta-analysis (Comte et al., 2013) pointed out heterogeneous responses of freshwater fish species to ongoing and anticipated climate change: some species are (or are predicted to be) responding positively, others are not. In this regard, the meta-analysis highlighted that cold-water species displayed reductions in their spatial distributions or distributional shifts to higher altitude, whereas cool and warm-water species showed distributional range expansions (Comte et al., 2013). Note, however, that the range expansion observed or predicted for some species is physically constrained. It can occur only within river basins that are already occupied; without human intervention, inhabited river basins will remain inhabited. For the species responding to climate change with a decrease in their area of occupancy, a higher risk of being extirpated from some river basins is expected.

To fully understand the dynamics of population loss, it is essential to distinguish between deterministic and stochastic extinctions (Lande, 1993), as the latter can lead to delayed extinctions even after the climate has stabilized. Deterministic extinctions occur when environmental conditions are no longer favorable in the long term, resulting in a negative population growth rate, making population extinction ineluctable (Figure 1). In the context of climate change, deterministic extinctions happen when climatic conditions become unfavorable across the entire potential area of occupancy of a population. Stochastic extinctions, on the other hand, arise from temporal variability in population size caused by environmental stochasticity (Caughley, 1994; Lande, 1993). In such cases, an extinction can occur even if the population occupies an overall favorable habitat where its expected growth rate is positive (Lande 1993). Small populations are particularly vulnerable to stochastic extinctions (Caughley, 1994), and in conservation biology, a population is considered non-viable if its size falls below a threshold known as the minimum viable population, MVP, (Shaffer, 1981). Climate change, by reducing the size of suitable habitats, may transform a viable population into a non-viable one, which is likely to go extinct in the short term. Predicting deterministic extinctions is straightforward if future habitat suitability can be modelled under a specific climate change scenario, using a species distribution model, for instance (Comte et al., 2016; Conti et al., 2014). If, in a given river basin and for a particular species, the predicted suitable area is null, a deterministic extinction will inevitably occur. Predicting which populations will become non-viable, however, requires an extinction model that translates population size into extinction probability and assesses whether the population size falls below the MVP (Figure 1). Since the sizes of many freshwater fish populations are unknown, even the simplest demographic models used in population viability analyses are often not feasible. To overcome this issue, area of occupancy is used in this study as a surrogate for population size. The relationship between population size and area of occupancy is a well-documented macroecological pattern, and occupancy data are more readily available than densities. Moreover, area of occupancy is relatively easy to model and predict future under a given climate change scenario by feeding a species distribution model (SDM) with presence/absence data. Finally, it has been shown that a model based on area of occupancy was a good predictor of population extinction rates observed during the Holocene in a system of French coastal rivers that became isolated due to rising sea level.

**Figure 1.**
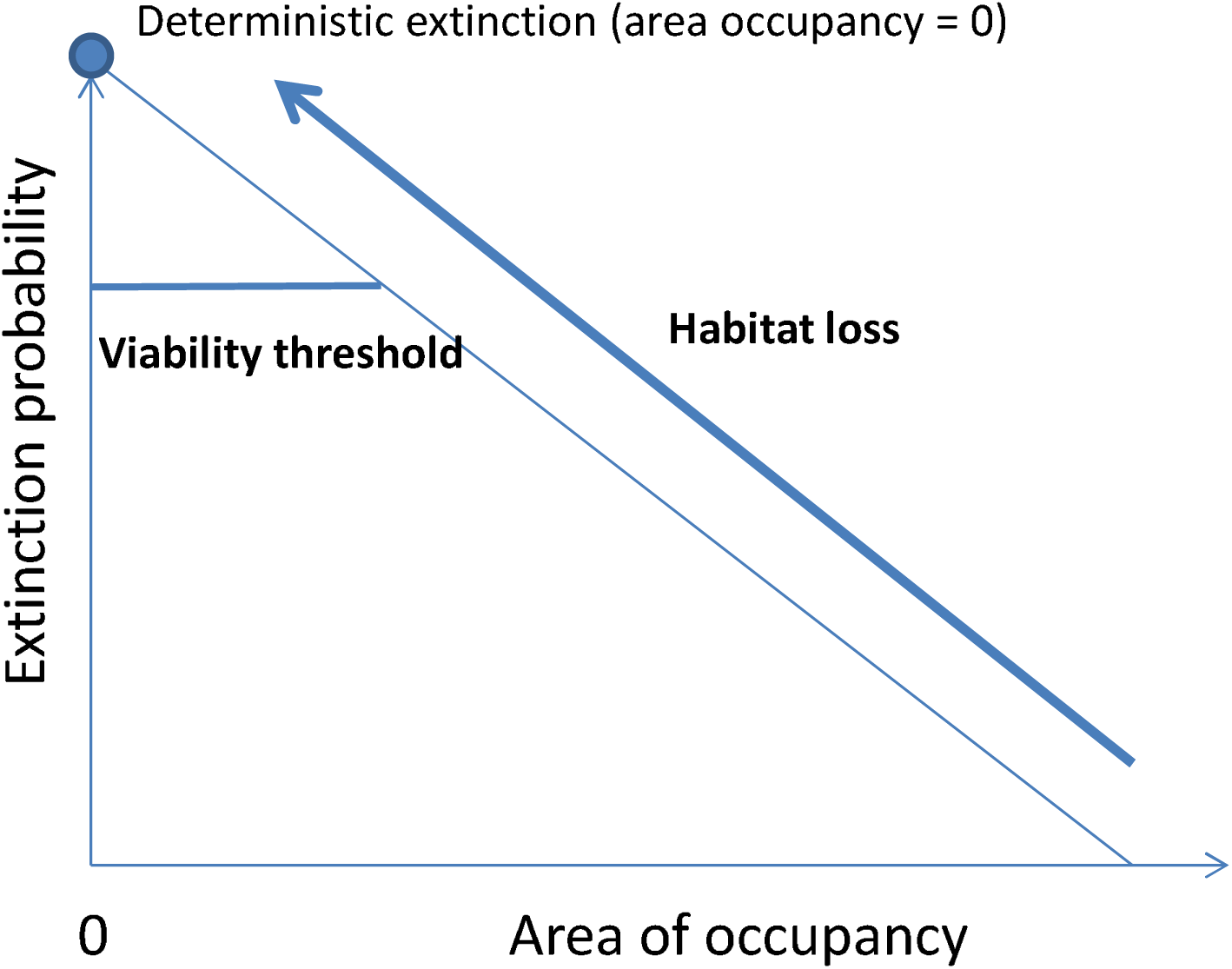
Conceptual framework of the study, defining deterministic extinctions. So as to simplify, we used linear relation between extinction and area of occupancy.

In this study, a former extinction model (Bellard & Hugueny, 2020; Hugueny et al., 2011) has been improved and re-parametrized so that it can be coupled with a SDM to predict the extinction rate of a population based on the area it is projected to occupy under future climatic conditions. By doing so, it allows for the assessment of the population’s viability status using area of occupancy as a surrogate for population size (Gaston & Fuller, 2009; He & Gaston, 2000) unknown for most freshwater fish species. This approach, which pairs SDM with EAR, is applied to the 23 most common freshwater fish species found across French territory to predict the effects of climate change on population extinctions. France is particularly suited for that purpose, as fish communities have been monitored extensively across the country since the 1995s, providing high-quality data for feeding presence-absence SDMs and calibrating EARs.

First, using current climatic conditions and presence/absence data from 1729 sites, we fitted Species Distribution Models (SDMs) to 23 freshwater fish species across the French territory to estimate their current and future distribution ranges (under the RCP 8.5 scenario for the year 2070), integrating both climatic but also geomorphological variables to estimate habitat distribution. Second, assuming no major climatic changes occurred during the Holocene, we used SDMs results for 15 species of these species in 18 French coastal basins, which became isolated from each other about 8500 years ago due to rising sea level. We then calibrated an empirical model to estimate population extinction rates as a function of area of occupancy at the time of isolation and the river basin area (extinction-area of occupancy model).

Third, using SDMs results for each river basin, we identified populations predicted to be extirpated due to complete loss of suitable habitat under future conditions (deterministic extinction) and compared these results to the value expected if no climate change had occurred (as predicted by the extinction-area occupancy model).

Fourth, in basins where species were predicted to persist, we applied the extinction-area of occupancy model to assess the extent to which climate-driven changes in occupancy have compromised the long-term persistence of populations. We inferred non-viable populations based on the minimum population viability threshold under current and future geo-climatic conditions. Finally, we tested whether cold-water species are more vulnerable to climate change by determining whether their local populations are more likely to shrink, be extirpated due to habitat loss, or become non-viable.

## Material and Methods

### Study area

This study was conducted on 162 basins across France, with significant differences in area between large basins (Loire, Rhône, Seine, Garonne) and smaller coastal catchments. The hydrological network studied comprised a total of 112,427 river segments, extracted from the RHT database (Pella et al., 2012), which served as a basis for projecting the SDMs models.

### Fish occurrences

Fish occurrence data were provided by the French Office for Biodiversity (OFB) at the national scale across France. We worked with data from 1729 sites sampled at least twice between 1995 and 2005 using an electrofishing protocol, with individuals caught between May and October each year. We focused on 25 of the most common species, belonging to eight families. The Cyprinidae family was the most diverse, with 15 species, while the other families included Esocidae, Salmonidae, Lotidae, Cottidae, Nemacheilidae, Cobitidae, and Percidae. For each site, we used yearly sampling records. For each species, we extracted the mean body length from the Fishbase database and spawning thermal preferences from Poulet et al., 2011, values which will be used in the extinction model.

### Current and future climatic and geo-morphological data

First, we used four geomorphological variables extracted from the RHT database: slope (‰), distance to the source (km), surface area of the drainage basin (km^2^) and river width (m) at the site. To eliminate co-linearity among the last three variables, which reflect the position along the upstream-downstream gradient, we conducted a principal component analysis (PCA) and the first axis, accounting for 90% of the variability, was kept as a synthetic variable describing the upstream–downstream gradient.

We extracted six climatic variables from the WorldClim database (Hijmans et al., 2005, version1.4) with a 30 arc-second resolution grid: mean temperature of coldest three months (BIO11), mean temperature of warmest three months (BIO10), temperature seasonality (BIO4), precipitation of the wettest three months (BIO16), precipitation of the driest three months (BIO17) and precipitation seasonality (BIO15). Future climate conditions were also extracted for each of these variables for the 2070s period, as an average forecasted over the period of 2061-2080. For this period, we used the CNRM (French national center of meteorological research) General Circulation Model (GCM) and focused on the most critical greenhouse emissions scenario (GES) (RCP 85).

### Species distribution models

The dataset of 1729 sites was split into a calibration (70 %) and a validation (30 %) dataset and this splitting was repeated 100 times. For each run, we computed 5 SDMs: generalized additive models (GAM), generalized boosted tree (GBM), random forests (RF), multivariate adaptive regression splines (MARS), and classification tree analysis (CTA). The predictions from the 5 models were averaged for current conditions using a consensus method based on the average values of the ensemble modeling predictions. and we assessed the predictive performances of the five statistical models and the “consensus model” using AUC. Then, we converted habitat suitability values (i.e., predictions from the consensus model) into occurrence probabilities by fitting a logistic relation between observed occurrences and habitat suitability values, and extracting the predictions from this relation as probabilities of occurrence. Finally, these models were also used to project future potential habitat suitability converted into occurrences probabilities as previously done across France, under the hypothesis of free dispersal and only for basins where the species was currently present.

### The extinction-area of occupancy model

The model described below is slightly modified from the one proposed by Bellard & Hugueny (2020). We modelled *e_ij_*, the probability of annual extinction of species *i* in river basin *j*, as a function of *L_i_*, the body length of species *i* (in cm), *p_ij_*, the proportion of occupied sites (occupancy) of species *i* in river *j*, and *A_j_*, the drainage area of river *j* (in km^2^) according to the following equation (Model 1):

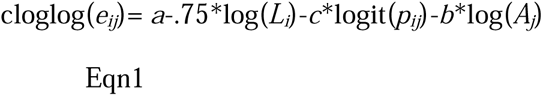

with cloglog(*e_ij_*)=log[-log(1-*e_ij_*)] and logit(*p_ij_*)=log[*p_ij_*/(1-*p_ij_*)].

This equation is based on several assumptions stemming from theoretical considerations or well-established empirical relationships. If environmental stochasticity is the primary driver of population fluctuations, then *e_ij_* is expected to follow a negative power function of *N_ij_* (Lande, 1993), the initial population size of species *i* in river *j*:

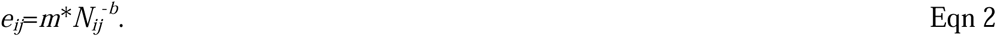

If only one individual is present, its death signifies the end of the population, and *m* is equivalent to the mortality rate. Note that it is assumed that neither parthenogenesis nor internal fertilization occurs, so that a single female cannot reestablish the population, as is the case for all the species considered in the present study. According to the metabolic theory of ecology (Brown et al., 2004), which states that mortality rate scales with body mass or body length with an exponent of -0.75 respectively, we modelled *m* as follows

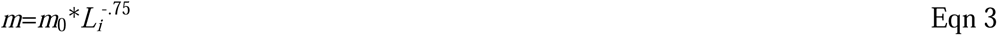

To avoid over-parameterization, we assumed that *b* was constant among species. Since its value cannot be anticipated based on theoretical works, it was treated as a free parameter.

In a second step, we assumed a positive relationship between the average population density per site for species *i* in river *j*, *d_ij_*, and the fraction of sites occupied, *p_ij_*. The positive correlation beween *d* and *p* is a well-documented empirical pattern, and several functions have been proposed in the literature to describe it (He & Gaston, 2000). Here, after preliminary analyses, we adopted the logistic function proposed by Hanski & Gyllenberg, 1997:

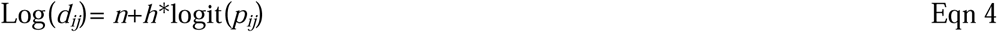

For the sake of simplicity, no interspecific variability or variability among river basins in the parameters *n* and *h* has been considered.

Therefore population size of species *i* in river *j* is expressed as

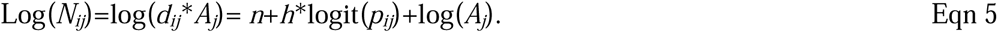

Combining Eqn 2, 3 and 5 and aggregating parameters when necessary leads to the right-hand side of Eqn 1. The left-hand side should be log(*e_ij_*), but the log function does not constrain *e_ij_* to be less or equal to one, which could be problematic for iterative parameter estimation. For small *e*, cloglog(*e*) is very close to log(*e*) and always returns *e*<1, so Eqn 5 preserves the expected relationship between *e* and *N* (Eqn 2) and enables *e* to behave as a probability during the estimation procedure.

### Fitting the extinction model

To estimate the parameters of Eqn 1, we took advantage of a natural fragmentation experiment allowing us to estimate the extinction rates of fish populations isolated in coastal rivers since the end of the Pleistocene. Bellard & Hugueny, 2020 and Hugueny et al., 2011 provided a detailed description of this system, which is briefly synthesized here. During Pleistocene sea-level low stands, the coastal rivers now discharging into the English Channel, an arm of the Atlantic Ocean separating France from England, were interconnected and formed a single river basin, the Channel River (Ménot et al., 2006). These river basins became isolated about 8,500 years ago due to rising sea level. (Hugueny et al., 2011) selected 18 French rivers that formerly belonged to the Channel River and used information about the distribution of 15 fish species to study extinction patterns. The same set of rivers and species is considered in the present study.

The probability of observing species *i* in coastal river *j*, *t* years after river isolation, *pa_ij_*, is given by (Bellard & Hugueny, 2020):

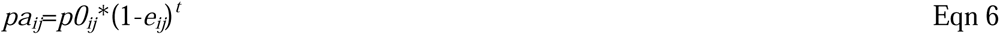

where *p0_ij_* is the probability of presence of species *i* in river *j* at the time of isolation, and *e_ij_* is given by Eqn 1. The time elapsed since isolation, *t*, is set to 8,500 years. The observed current presence-absence matrix (*pa_ij_*) is provided by (Hugueny et al., 2011). Whether or not a given species was present in a specific river at the time of its isolation from the palaeo-river network is unknown and needs to be estimated. Based on the presence-absence data of species within 37 tributaries of the Seine River, used as a reference condition for non-isolated river basins, an incidence function for each species’ probability of presence in a tributary as a function of its surface area was established in previous works (Bellard & Hugueny, 2020; Hugueny et al., 2011) and re-used here to predict *p0_ij_* knowing *A_j_*.

Assuming no major climatic changes during the Holocene, we used the SDMs fitted to each species to predict *p_ij_* in Eqn 1. Although warm phases (e. g. Roman Climatic Optimum) and cold phases (e.g. Little Ice Age) alternated during the Holocene, palynological studies ((Davis et al., 2003; Renssen & Isarin, 2001) suggest that average thermal conditions have been relatively close to pre-industrial values in the region of interest for most of the past 8.500 years.

The parameters of Eqn 1 were estimated by maximizing the likelihood of observing *pa_ij_*. For this purpose, we used the “nlminb” function (package “stats”) in the R environment, assuming a Bernoulli distribution for *pa_ij_*. We also considered models that excluded occupancy (Model 2) or body size (Model 3) to verify the importance of these variables in predicting species occurrences. Models were compared using the Akaike Information Criterion. The quality of fit of Model 1 was visualized by plotting the number of basins currently occupied per species against the number predicted (by summing, for each species, the predicted *pa_ij_*).

### Statistical analyses

For each species, we used the previously described extinction model to determine deterministic and stochastic extinction rates.

First, we calculated the percentage of river basins with deterministic extinction as the ratio between the number of basins exhibiting a probability of presence equal to zero under future climatic scenario (i.e., no suitable habitat remaining for the species) and the number of basins where habitat loss was projected between future and current distributions. Stochastic (or background) extinctions were determined using the extinction model based on the current probability of occurrence of the species, assuming habitat stability. We then calculated the ratio between the percentage of deterministic extinctions and the stochastic extinction rate to compare these two types of extinctions.

Second, we calculated a multiplication factor to test the effect of climate change. This factor corresponded to the ratio between the extinction rate under future climatic conditions (based on species’ probability of presence from SDMs using future climatic conditions) and the extinction rate under current climatic conditions.

Third, we assessed population viability for each species under both current and future climatic conditions. The Minimum Viable Population (MVP) is defined in this study as the population size corresponding to a 0.01 probability of extinction over a 40 generations period which for vertebrates corresponds to about 4000 individuals (Reed et al., 2003; Traill et al., 2007). Assuming a constant extinction rate over time, a population is therefore considered viable if its modelled annual extinction probability is lower than a threshold of 1-(.99)^(1/40*Tg), with Tg the generation length of the focal species.

## Results

### Calibration of extinction model

The estimated parameters for Model 1 are as follows: *a*=-1.603, *c*= 0.887, *b*=1.091. Both *c* and *b* were positive as expected. This model provided a very good fit (r=0.941) to the number of rivers occupied per species (Figure 2). Both occupancy and body size contributed significantly to Model 1, as indicated by the higher values of AIC values obtained for Model 2 (301.38, without occupancy) and Model 3 (183.56, without body size), compared to Model 1 (175.16).

**Figure 2.**
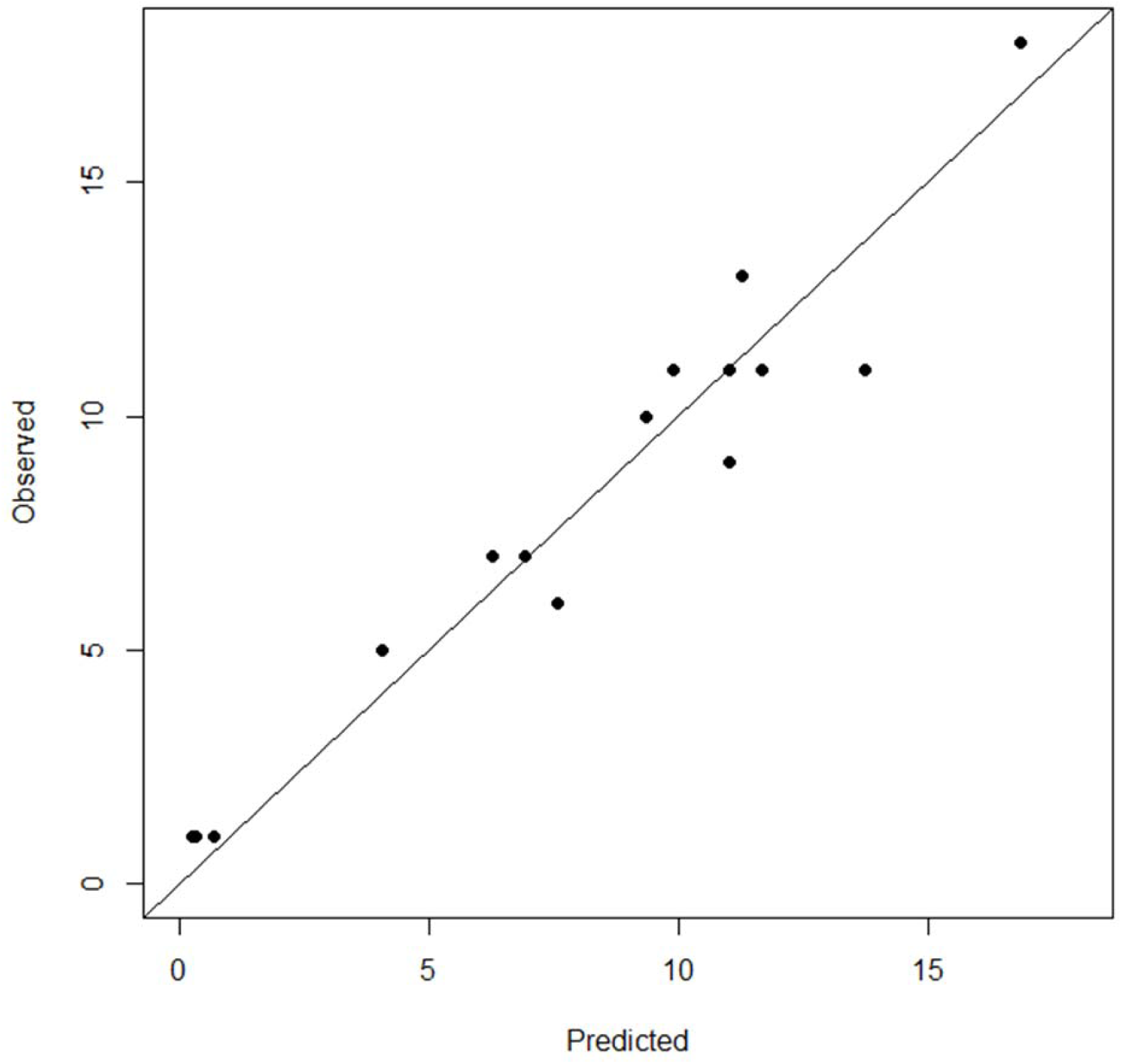
Number of river basins predicted to be occupied by the extinction-area of occupancy model (see text) vs. observed values. The line of perfect fit is shown. Each point represents a species.

### Species distribution modeling

Results showed a loss of distribution area, especially in northern France for some species such as *Cottus gobio*, *Perca fluviatilis, Esox lucius* and *Salmo trutta*. Small northern rivers were significantly impacted (Figure 3, Table1), especially for *C.gobio* and *S.trutta* species. In contrast, other species, particularly those with broader distributions such as barbel (*Barbus barbus*) or European minnow (*Phoxinus phoxinus*), exhibited an increasing presence across France under future conditions (Figures 1 and S1, Table 1). Stability in the distribution area was observed for two currently widely distributed species: gudgeon (*Gobio gobio*), hotu (*Chondrostoma nasus*) and loach (*Barbatula barbatula*), with a weak increase for *G.gobio* (Figure S1, Table 1).

**Figure 3.**
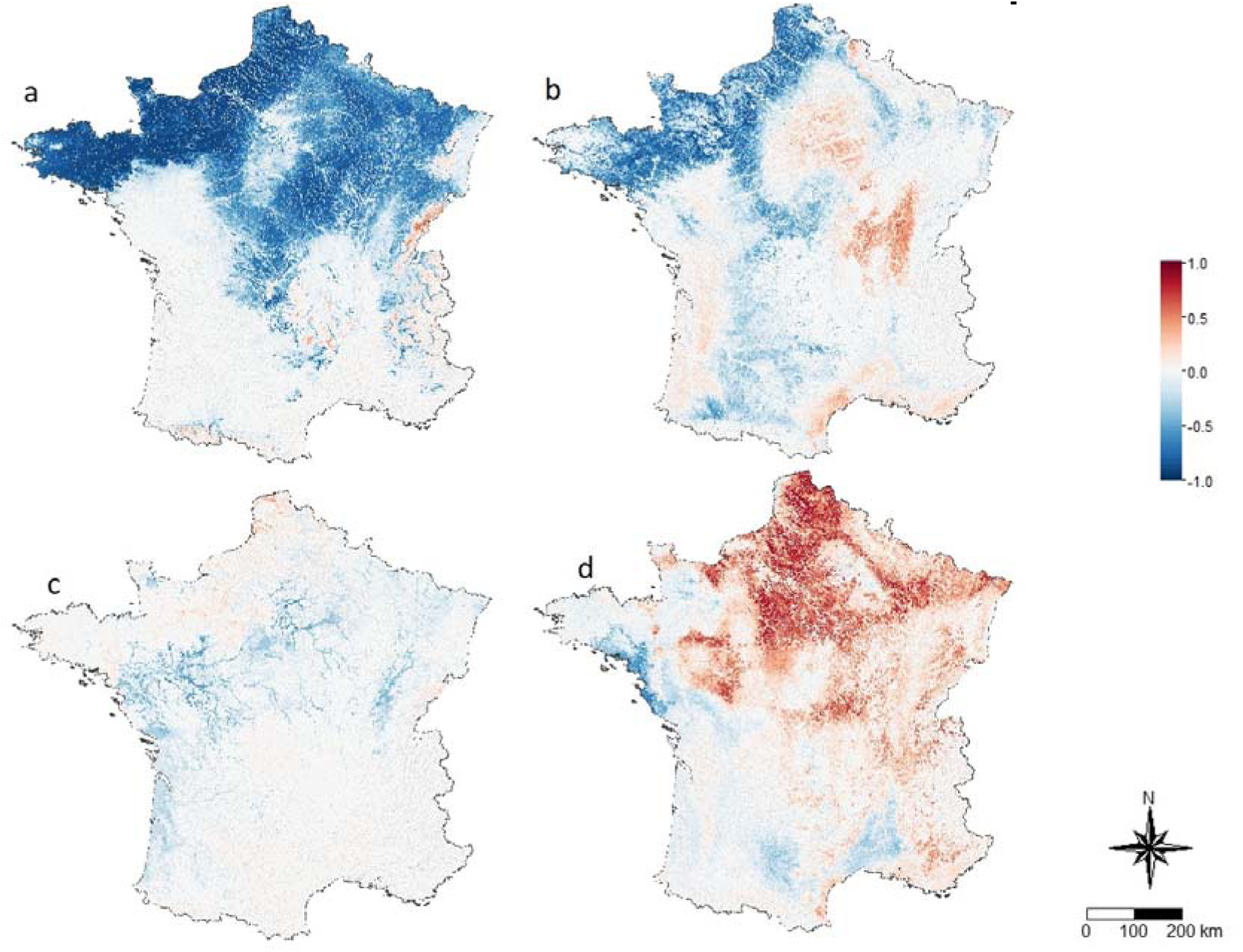
Changes in suitability index for four species between future and current conditions for a) bullhead (*Cottus gobio*), b) common trout (*Salmo trutta*), c) pike (*Esox Lucius*) and d) european minnow (*Phoxinus phoxinus*) (values of suitability habitat are presented for other species in Appendix S1). Positive values (in red) indicate an increase of suitability for the species, while negative ones (in blue) indicate a decrease of suitability in the future.

**Table 1.** Modelled sum of current and future occurrences based on species distribution modeling in basins where the species is currently present. The percentage of lost/gain habitat is the ratio between future and current occurrences.

| Latin species name | Sum of current occurrences | Sum of future occurrences | Species range change (%) |
| --- | --- | --- | --- |
| *Abramis brama* | 7080.19 | 5171.92 | -26.9 |
| *Alburnus alburnus* | 10782.08 | 14080.90 | +30.6 |
| *Barbatula barbatula* | 52275.45 | 51867.02 | -0.8 |
| *Barbus barbus* | 10603.89 | 18246.82 | +72.1 |
| *Barbus meridionalis* | 3401.36 | 10399.23 | +205.7 |
| *Blicca bjoerkna* | 3711.72 | 1048.71 | -71.7 |
| *Chondrostoma nasus* | 3174.59 | 3266.65 | +2.9 |
| *Cobitis taenia* | 418.14 | 35.52 | -91.5 |
| *Cottus gobio* | 38757.52 | 8381.05 | -78.4 |
| *Cyprinus carpio* | 6036.05 | 3784.57 | -37.3 |
| *Esox Lucius* | 10318.98 | 5707.70 | -44.7 |
| *Gobio gobio* | 43828.35 | 50485.79 | +15.2 |
| *Leuciscus leuciscus* | 9710.67 | 7583.36 | -21.9 |
| *Lota lota* | 275.17 | 60.59 | -78 |
| *Perca fluviatilis* | 23570.89 | 13372.52 | -43.3 |
| *Phoxinus phoxinus* | 46192.17 | 57026.92 | +23.4 |
| *Rhodeus sericeus* | 4493.42 | 2350.60 | -47.7 |
| *Rutilus rutilus* | 33416 | 28621.51 | -14.3 |
| *Salmo trutta* | 61509.73 | 51093.15 | -16.9 |
| *Scardinius erythrophthalmus* | 9780.99 | 4288.91 | -56.1 |
| *Squalius cephalus* | 36316.22 | 52807.10 | +45.4 |
| *Thymallus thymallus* | 1159.75 | 486.43 | -58.1 |
| *Tinca tinca* | 10014.73 | 4835.15 | -51.7 |

### Deterministic vs stochastic extinctions

As expected, deterministic extinctions due to habitat loss from climate change occurred for some populations (Table 2). However, half of the studied species (11 out of 23) exhibited no expected deterministic extinctions (e.g., *Squalius cephalus* and *G. gobio*) (Table 2). For some cold-water species such as bullhead (*C. gobio*), the number of modelled deterministic extinctions was more than 10 times higher than would have been expected in the absence of climate change (Table 2). Overall, among declining populations, about 3% were predicted to be extinct by 2070 due to complete loss of suitable habitat. The number of deterministic extinctions was, on average, 7 times higher than stochastic extinctions under the assumption of no change in suitable habitat area. These deterministic extinctions were mainly located in the small western coastal rivers of the English Channel and Brittany for several species (Figure 4).

**Table 2.**
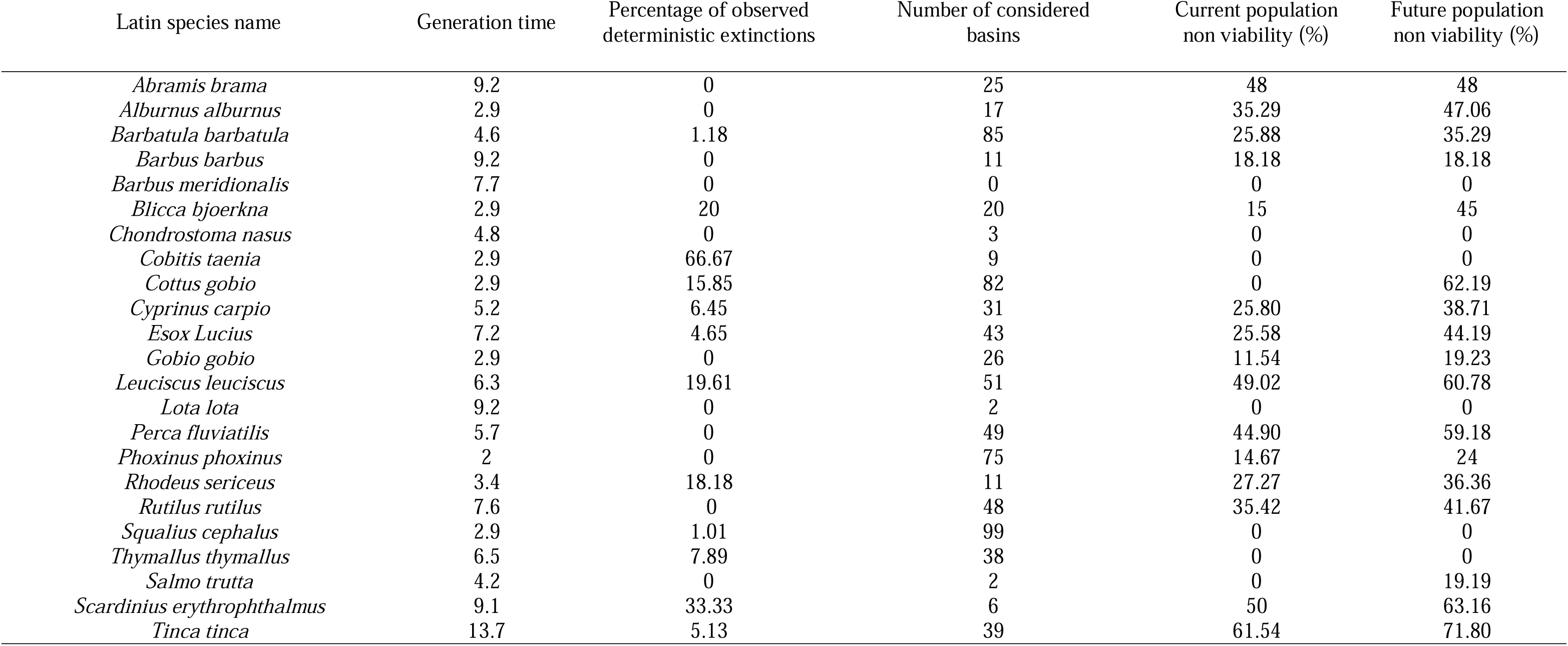
Percentage of deterministic extinctions per species (river catchments where the species is present) and percentage of non-viability of populations for each species under current and future climatic conditions for one rate of Minimum Population Viability (MVP) and for each freshwater species in basins where the species is present and a loss of habitat is observed. The percentage of observed deterministic extinctions is equal to the ratio between the number of basins exhibiting a value of area equal to zero under future climatic conditions on the number of basins where a loss of area is observed for the species. The background or stochastic extinction rate corresponds to the sum of extinction rate values in 2070 across considered basins (results of extinction model without climate change).

| Latin species name | Generation time | Percentage of observed deterministic extinctions | Number of considered basins | Current population non viability (%) | Future population non viability (%) |
| --- | --- | --- | --- | --- | --- |
| *Abramis brama* | 9.2 | 0 | 25 | 48 | 48 |
| *Alburnus alburnus* | 2.9 | 0 | 17 | 35.29 | 47.06 |
| *Barbatula barbatula* | 4.6 | 1.18 | 85 | 25.88 | 35.29 |
| *Barbus barbus* | 9.2 | 0 | 11 | 18.18 | 18.18 |
| *Barbus meridionalis* | 7.7 | 0 | 0 | 0 | 0 |
| *Blicca bjoerkna* | 2.9 | 20 | 20 | 15 | 45 |
| *Chondrostoma nasus* | 4.8 | 0 | 3 | 0 | 0 |
| *Cobitis taenia* | 2.9 | 66.67 | 9 | 0 | 0 |
| *Cottus gobio* | 2.9 | 15.85 | 82 | 0 | 62.19 |
| *Cyprinus carpio* | 5.2 | 6.45 | 31 | 25.80 | 38.71 |
| *Esox Lucius* | 7.2 | 4.65 | 43 | 25.58 | 44.19 |
| *Gobio gobio* | 2.9 | 0 | 26 | 11.54 | 19.23 |
| *Leuciscus leuciscus* | 6.3 | 19.61 | 51 | 49.02 | 60.78 |
| *Lota lota* | 9.2 | 0 | 2 | 0 | 0 |
| *Perca fluviatilis* | 5.7 | 0 | 49 | 44.90 | 59.18 |
| *Phoxinus phoxinus* | 2 | 0 | 75 | 14.67 | 24 |
| *Rhodeus sericeus* | 3.4 | 18.18 | 11 | 27.27 | 36.36 |
| *Rutilus rutilus* | 7.6 | 0 | 48 | 35.42 | 41.67 |
| *Squalius cephalus* | 2.9 | 1.01 | 99 | 0 | 0 |
| *Thymallus thymallus* | 6.5 | 7.89 | 38 | 0 | 0 |
| *Salmo trutta* | 4.2 | 0 | 2 | 0 | 19.19 |
| *Scardinius erythrophthalmus* | 9.1 | 33.33 | 6 | 50 | 63.16 |
| *Tinca tinca* | 13.7 | 5.13 | 39 | 61.54 | 71.80 |

**Figure 4.**
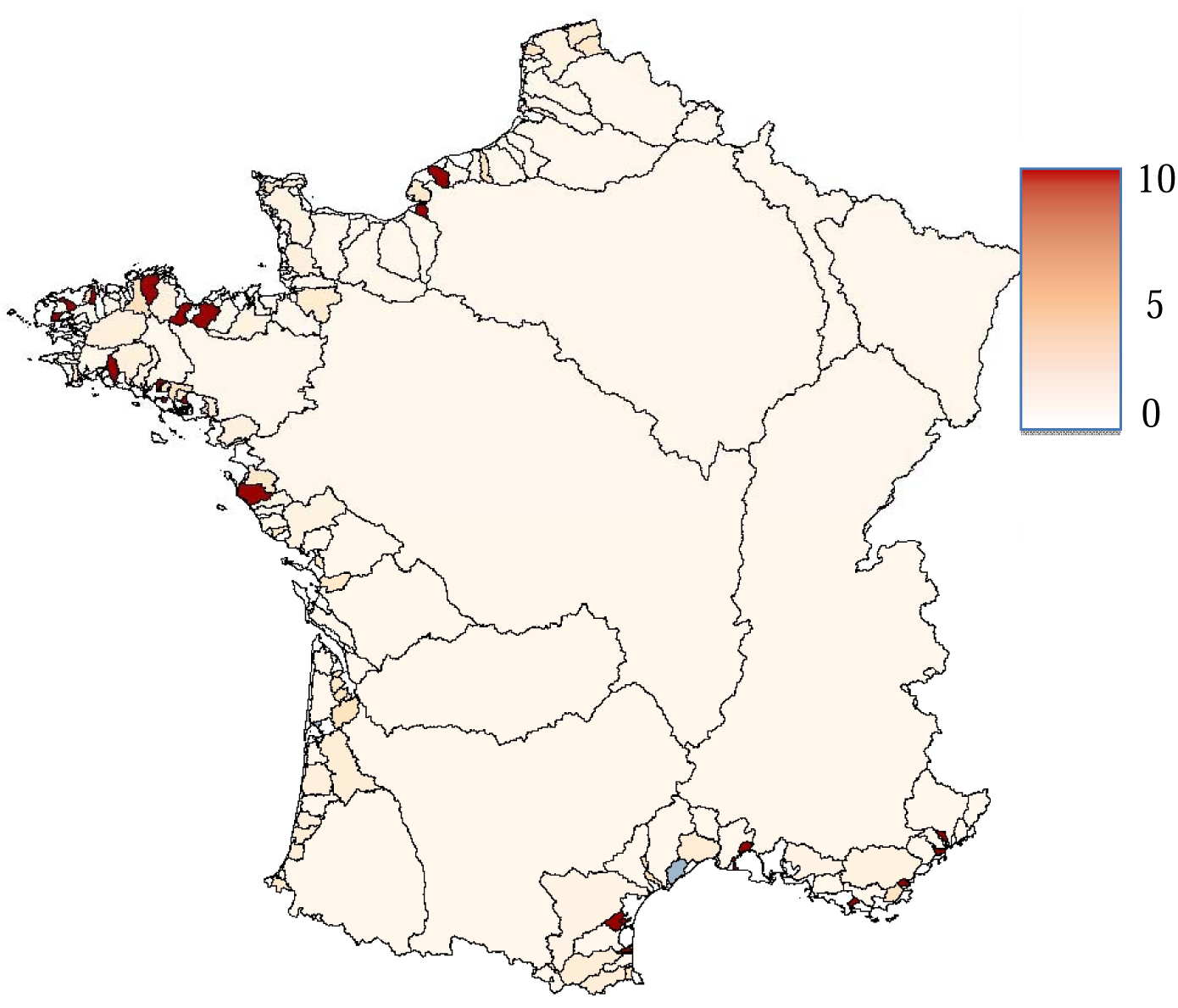
Spatial patterns in the percentage of fish species displaying deterministic extinctions under future climatic conditions.

### Spatial patterns of extinctions and impacts of climate change

Ratios between extinction rates under future climatic conditions and extinction rates under current conditions were very high for *C. gobio* and *S. trutta*, particularly in small western coastal basins of the English Channel and Brittany, with values ranging from 0 to 140 for *C. gobio* and from 0 to 50 for *Salmo salar*. Strong values were also observed for pike (*E. lucius*) and minnow (*P. phoxinus*) in the western part of France (Figure 5). For other species, large basins like the Rhône River basin were also impacted for some species (*Rhodeus sericeus*, *Abramis brama*, *Cyprinus carpio* and *Scardinius erythrophtalmus*) (Figure S2), though small coastal basins were the most impacted for species such as *A. alburnus*, *Barbus meridionalis*, *Barbus barbus* and *Barbatula barbatula*.

**Figure 5.**
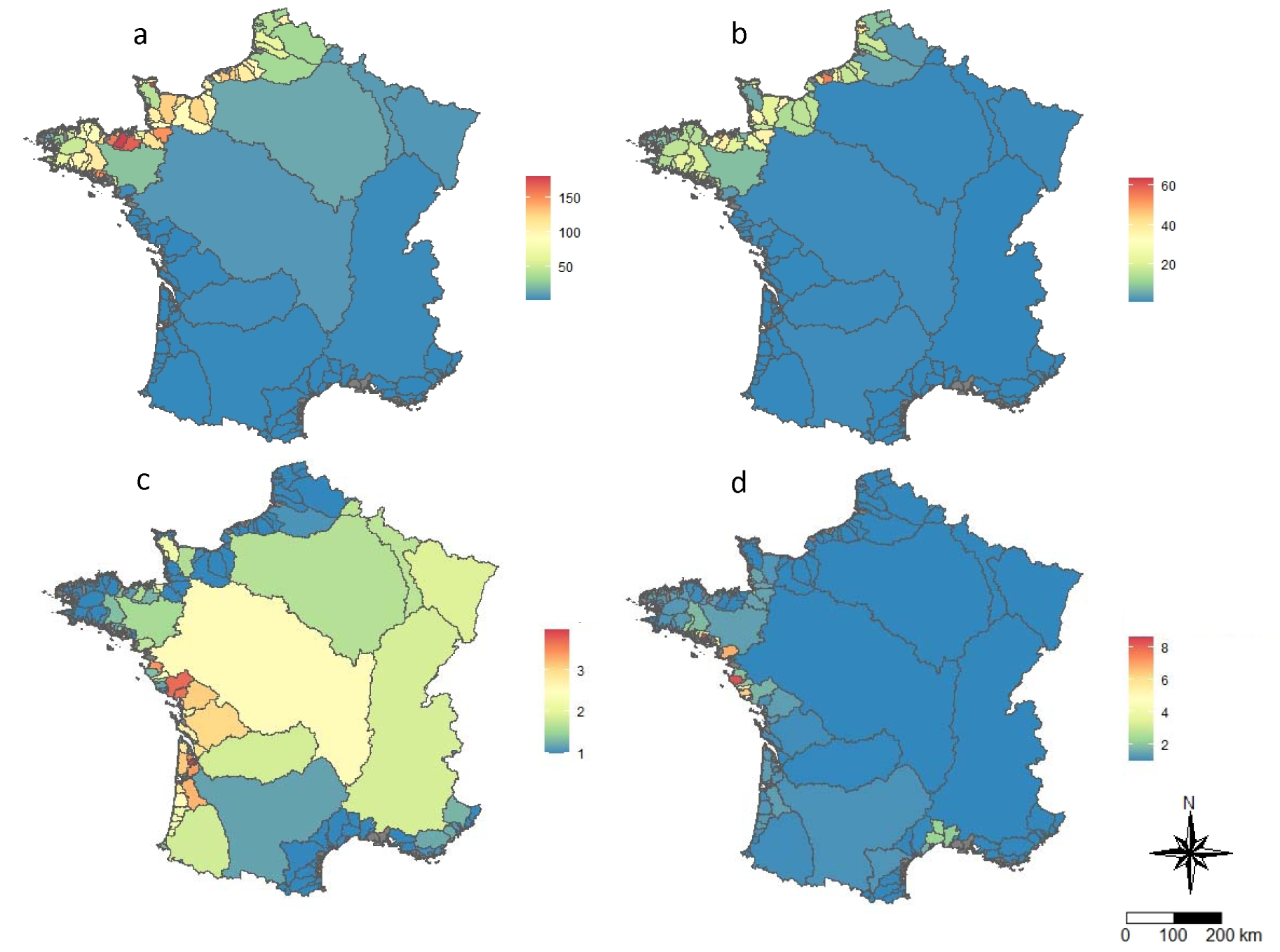
Ratio between stochastic extinction rate under future climatic conditions and extinction rate under current conditions for each river basin for a) bullhead (*Cottus gobio*), b) common trout (*Salmo trutta*), c) Pike (*Esox lucius*) and d) european minnow (*Phoxinus phoxinus*) (other species are presented in supplementary material).

### Viability of populations

For a restrictive MVP (1% in 40 generations) under current climatic conditions, high rates of non-viability were observed for *A. alburnus*, *R. sericeus*, *A. brama*, *E. lucius*, *C. carpio*, *Rutilus rutilus*, *B. barbatula*, *Perca fluviatilis*, *Tinca tinca* and *Leuciscus leuciscus*. Significant increases in the percentage of non-viable populations from current to future climatic conditions were also observed for *Blicca bjoerkna*, *E. lucius*, *C. gobio*, *S. trutta* and *L. leuciscus* (Table 2). Overall, there is an increase of ratio of non-viable populations in the future. Only *A. brama* and *B. barbus* remained totally stable, with no decreases observed for any species. The average current rate of non-viable populations is 21.2 % across all species, while the average future rate is 31.9%.

### Extinction risk and spawning temperature

Considering three metrics of extinction risk, a concave relationship was observed with the spawning temperature of the species (Figure 6), with higher extinction risk for cold (5-10°C) and warm (20-25°C) water species. According to AIC values, the fraction per species of populations that are predicted to loss habitat is better fitted by a logistic model with temperature modelled by a second order polynomial rather than by a linear trend (delta AIC= - 59.15). Regarding the fraction of populations predicted to vanish because of complete habitat loss, a concave relationship is also the best model (delta AIC= -27.67), as it is for the fraction of viable populations predicted to turn into un-viable ones ((delta AIC= -11.63)

**Figure 6.**
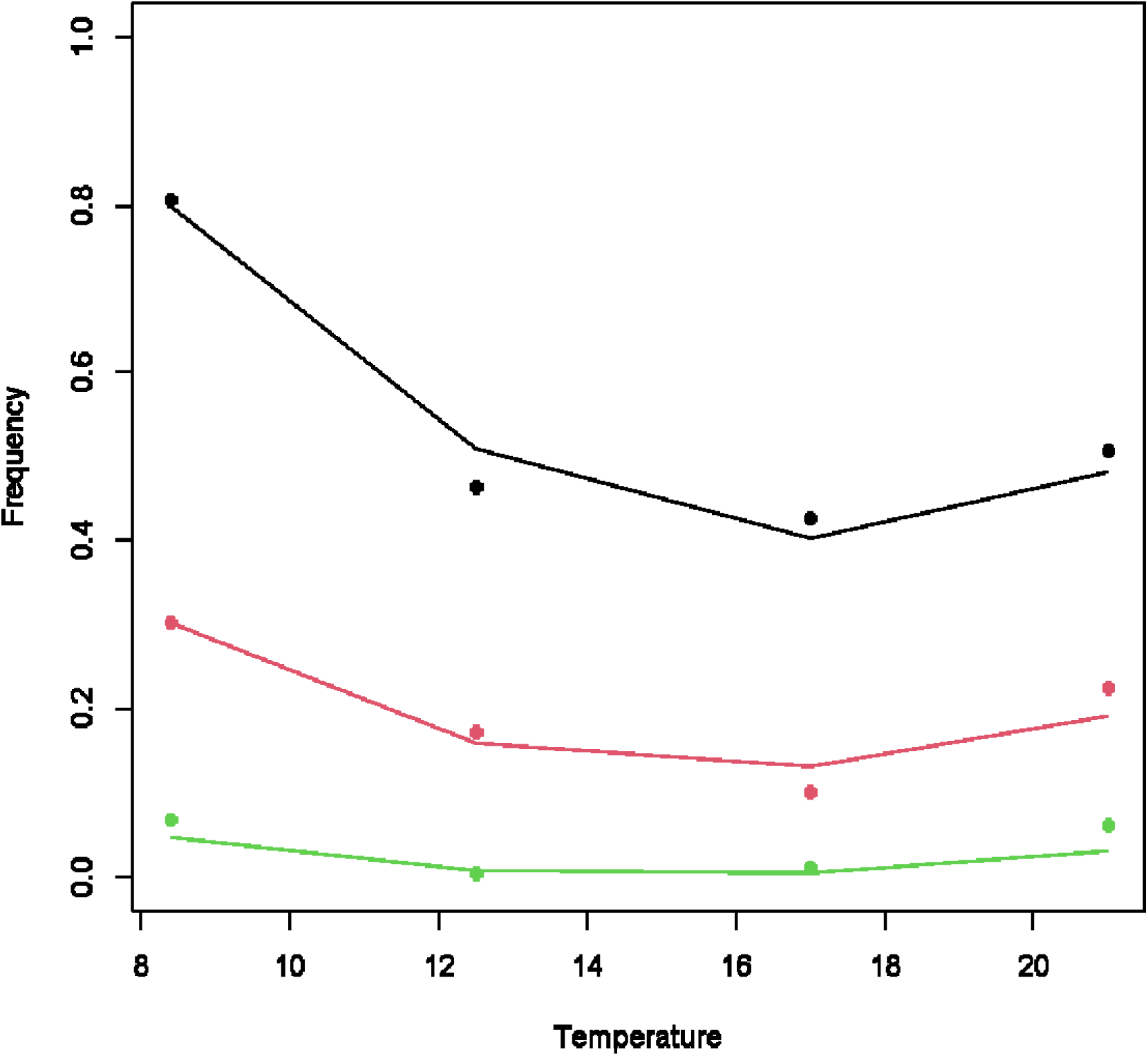
Curvilinear relationships relating three metrics of extinction risk and the spawning temperature. Species have been grouped in accordance to their spawning temperature, [5-10[, [10-15[, [15-20[ and [20-25[ °C, and each group is plotted at its average temperature. Dots and lines represent observed values and predicted values from quadratic logistic models, respectively, averaged per group in each case. Black indicates the frequency of populations with habitat loss, red indicates the ratio of populations with deterministic extinctions over the number of populations experiencing habitat loss, and green indicates the frequency of viable populations turned into non-viable ones after habitat loss.

## Discussion

Our results showed a significant predicted effect of climate change on three indices of extinction risk, based respectively on the decline in the area of occupancy, extinctions due to lack of suitable habitat, and long-term population viability. Regardless of the index used, populations of cold-water species were the most threatened with extinction, followed by warm-water, and lastly, those with intermediate thermal preferences. These extinction patterns were spatially structured and could affect the viability of some populations.

### Expansion or contraction of distribution area according to species (SDM results)

Cold- and cool-water species, such as bullhead (*C. gobio*), trout (*S. trutta*), grayling (*Thymallus thymallus*), and pike (*E. Lucius*), are particularly negatively impacted by climate change, with a significant percentage of populations predicted to decline. In contrast, for a few warm-water species, our results predicted little to no population decline, especially for species currently located in southern France (e.g., *B. meridionalis*) and a widely distributed species (*S. cephalus*). More broadly, our results align with studies highlighting the vulnerability of cold-water freshwater fishes (population decline, range contraction) to anticipated climate changes in temperate regions (Comte et al., 2013; Myers et al., 2017). A simple explanation for the predicted contraction of cold-water species’ ranges is that in low-elevation coastal basins, these species have limited space to shift their range upstream. However, our results suggest that the relationship between thermal niche and the likelihood of population decline is more complex: extreme warm-water species tend to have a slightly higher proportion of declining populations than species with more intermediate thermal requirements. One possible explanation is that warm-water species prefer large, downstream rivers, and their ability to shift upstream is constrained by unsuitable habitats. This emphasizes the importance of considering factors beyond climate variables when modeling future species distributions. Similar contrasted results in the distribution shifts of freshwater fish have been previously observed, with some species like *Gymnocephalus cernua, A. alburnus*, and *C. carpio* showing increased spatial distribution, while others species like *T. tinca* exhibited decreased distribution (Comte et al., 2014). Variables such as mean temperature and precipitation were notably important in predicting the distribution of fish species across France, although the responses to these variables depended on the species considered (Grenouillet et al., 2011). This result is in agreement with a previous study conducted on the same species and at the same spatial scale (Buisson et al., 2008), which highlighted the significance of thermal and upstream-downstream gradients in species distribution. Some studies have also shown that responses to environmental factors are species-dependent and are close to their physiological and thermal requirements (curvilinear relation) (Comte & Grenouillet, 2013).

### Spatial variation in deterministic extinctions among species

The interaction between habitat loss and climate change can be highly detrimental, with species in fragmented habitats suffering more from climate change than those in non fragmented habitats (Travis, 2003). Our results showed that 1/5 of species experienced more than 20 % deterministic population extinctions. Similarly, Thomas et al., 2004 predicted that by 2050, 8 to 19 % of species from various taxa (birds, mammals, amphibians, reptiles, plants) will face deterministic extinctions at a global scale. The rate of deterministic extinctions observed in our study was notably important for cold-water species such as bullhead (*C. gobio*) and trout (*S. trutta*), but also for some warm-water species like bream (*A. brama*). In butterflies, it has been shown that specialist species were more impacted by climate change than generalist ones at a local scale (Franzén & Johannesson, 2007). Conversely, for ubiquitous species such as gudgeon and chub, we observed lower rates of deterministic extinctions. Higher extinction rates were also observed within small coastal river basins in northwestern France, revealing that deterministic extinction rates were spatially structured. In these basins, where the upstream-downstream temperature gradient is low due to their small size and low elevation, species have limited ability to cope with warming by moving upstream. This limitation could explain the higher occurrence of predicted deterministic extinctions in such areas.

### Extinction model

Our extinction model assumes that species loss occurred in coastal rivers following their isolation (Hugueny et al., 2011), because in such isolated basins, extinctions could not be balanced by colonizations since strictly freshwater fish species cannot colonize neighbor basins through sea water (Dias et al., 2014; Hugueny, 1989b; Oberdorff et al., 1997). This hypothesis has been corroborated in another freshwater systems, such as lakes formerly connected to the Baltic Sea, where fish extinctions occurred after the lakes became isolated by post-glacial land uplift (Bellard et al., 2019). As observed in other insular habitats (Hugueny, 2017), extinction rates were negatively correlated with the surface area of the “islands” (river basins or lakes).

Our model differs from previous approaches by adopting a population-centered approach. By modelling the area of occupancy for each species in each river, we were able to focus on extinction rates at the population level. This approach revealed that the rates of deterministic extinctions, in relation to surface area and biological parameters (such as body size), increased for most species. We confirmed the hypothesis that the most impacted species exhibit specific thermal and physiological requirements. However, our models did not account for other disturbances, such as anthropogenic fragmentation within river basins caused by dams or pollution. These factors can also affect extinction rates and significantly increase species vulnerability, suggesting that the actual scenarios could be more pessimistic than those we predicted. A previous study has already shown that recent extinction rates, including anthropogenic threats, could be 130 times higher than extinction rates predicted solely by water availability loss in rivers (Tedesco et al., 2013).

### Climate-induced changes in species habitat suitability as an important magnifying factor for local extinction

Overall the number of endangered species and the impact of habitat loss on biodiversity are often underestimated (Hanski & Ovaskainen, 2002). In our study, climate change emerged as a critical driver of habitat loss. Of the 23 species considered, 10 showed a higher rate of extinction than expected based solely on demographic factors (stochastic extinctions). This indicates that climate change-induced extinction rates are significantly higher than stochastic extinction rates, depending on the species. Similar findings have been reported in freshwater fish regarding water availability loss at a global scale (Tedesco et al., 2013), revealing that the median extinction rate was 7% higher than the background extinction rate. The local extinctions modelled in this study are realistic and are currently being observed in various species and taxa. While these local extinctions are difficult to quantify in freshwater fish, they have been documented in plants (Nomoto & Alexander, 2021) and insects (Macedo et al., 2016), especially through altitudinal shifts.

### Viability of populations

Our results showed that the viability of fish populations under future climatic conditions could be severely impacted, particularly for certain species. This is especially true for thermal specialist species such as *C. gobio*, for which a tenfold increase in extinction risk is expected from current to future conditions. The effects of climate change and land-use changes have been demonstrated in several taxa, affecting various demographic parameters at the species level (Hadjou Belaid et al., 2018; Kissel et al., 2019; Selwood et al., 2015; Sletvold et al., 2013). Similar to Tedesco et al. (2013), our results do not indicate alarming predictions in riverine fishes in France, except for some cold-water species. The time frames involved may provide an opportunity to implement conservation measures. This study underscores the need to prioritize conservation efforts for specific species. The model developed here, combined with species distribution modeling -an increasingly common tool in rewilding (Jarvie & Svenning, 2018) could be highly beneficial for conservation management. It could help identify priority areas for reintroduction efforts, as well as areas where species are likely to persist given future climatic conditions.

## Acknowledgments

This work has been supported by the BiodivERsA project ODYSSEUS funded from the European Union’s Horizon 2020 research and innovation program. CRBE lab was supported by “Investissement d’Avenir” grants (CEBA, ref. ANR-10-LABX-0025; TULIP, ref. ANR-10-LABX-41).

