## Supplementary files for "Climate-driven habitat loss and natural fragmentation increase extinction risk and compromise population viability in freshwater fish"

Supplementary material

Figure S1. Changes in suitability index for species considered in the study between future and current conditions. Positive values (in red) indicate an increase of suitability for the species, while negative ones (in blue) indicate a decrease of suitability in the future.

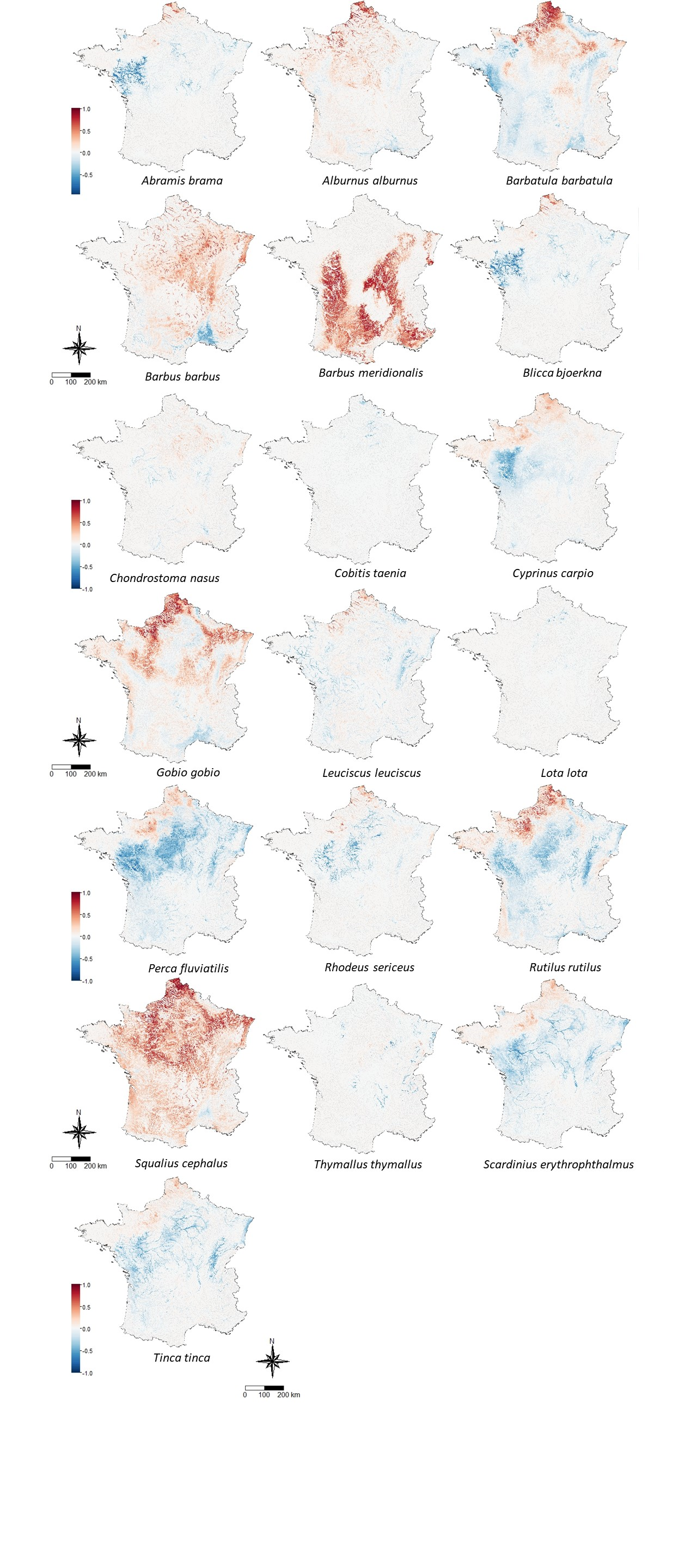

Figure S2: Ratio between future and current extinction rates for all studied species.

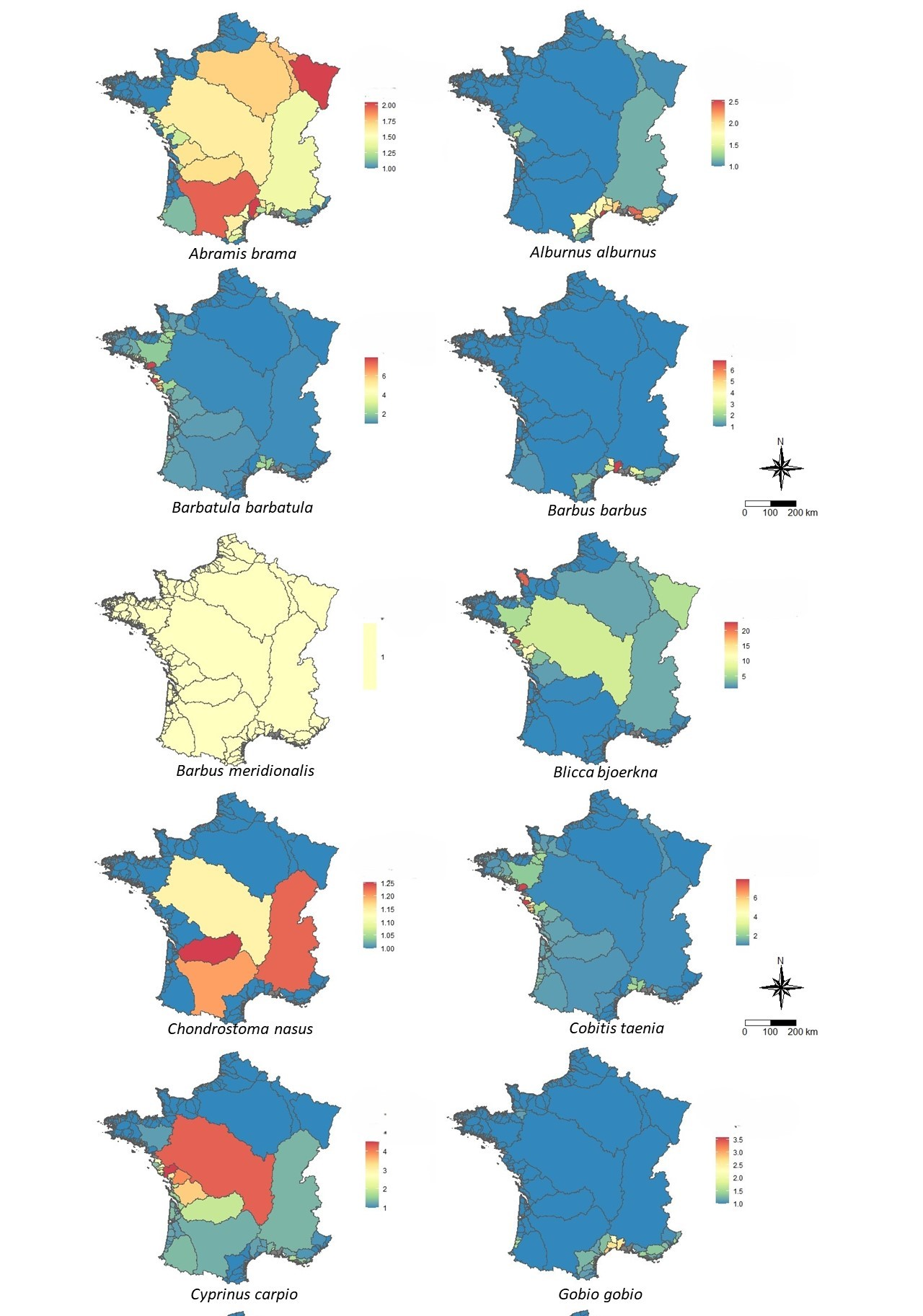

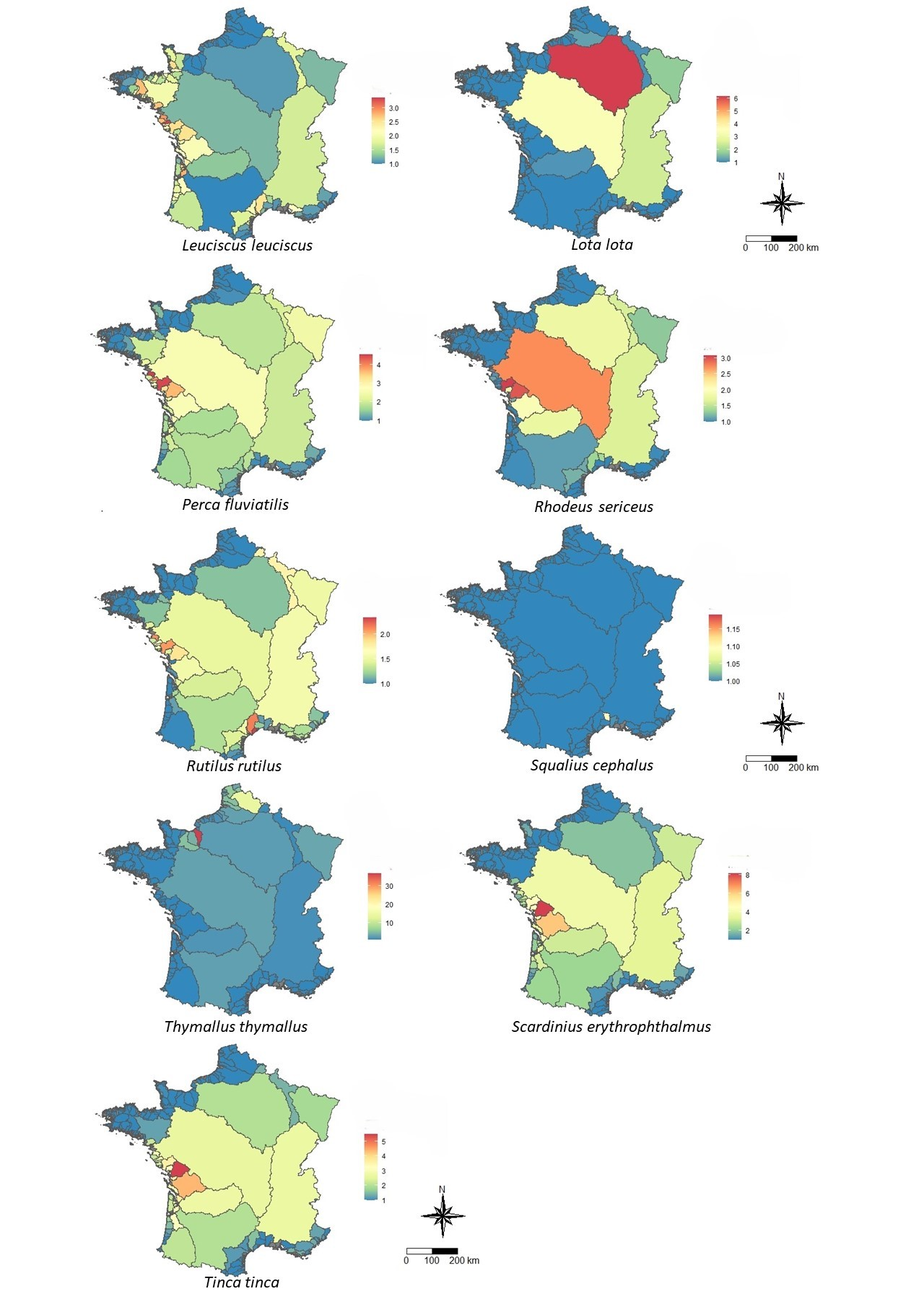
